# 4-HNE reduces phagocytosis through the expression of Synaptotagmin 1 in human monocyte-derived macrophages

**DOI:** 10.1101/2025.11.24.690103

**Authors:** Melina Ioannidis, Martijn den Ouden, Sietse J. Dijt, Danny Incarnato, Pieter Grijpstra, Joke Wortel, Matthijs Verhage, Sjors Maassen, Geert van den Bogaart

## Abstract

Reactive oxygen species (ROS) react with polyunsaturated fatty acids (PUFA) and generate the reactive aldehyde 4-hydroxynonenal (4-HNE). 4-HNE is a potent modulator of cell signaling, proliferation, and death. Our transcriptomics analysis revealed that upon treatment of human monocyte-derived macrophages with 4-HNE synaptotagmin-1 (SYT1), the main calcium sensor for neurotransmitter release, became the most strongly upregulated protein. This is surprising, as SYT1 expression is normally restricted to neurons and neuroendocrine cells. Using overexpression of SYT1 fused to a fluorescent reporter protein, we found SYT1 predominantly locates at the plasma membrane in macrophages. Based on this finding, and on the reported roles of other SYT forms in macrophages and other immune phagocytes, we tested the role of SYT1 in phagocytosis. Functional assays showed that SYT1 inhibited phagocytosis of pathogenic bacteria. Thus, our findings reveal an unexpected role of SYT1 in immune cells.

## Introduction

Macrophages are essential for innate cellular immunity. They are highly plastic and functionally diverse cells found in all tissues. Macrophages contribute to homeostasis, tissue repair, and immunity. They defend the host against pathogenic microorganisms, including bacteria, viruses, fungi, and parasites (Gordon, 2007; Murray & Wynn, 2011; Wynn et al., 2013). Macrophages express numerous surface receptors, intracellular mediators, and secretory molecules to recognize, engulf, and destroy pathogens (Taylor et al., 2005). Upon stimulation with microbial stimuli, such as lipopolysaccharide (LPS) of the outer cell membrane of gram-negative bacteria, macrophages develop a pro-inflammatory phenotype hallmarked by the expression of large amounts of pro-inflammatory mediators, including Tumor necrosis factor (TNF)-α, and Interleukin (IL)-6. Moreover, macrophages produce large amounts of reactive oxygen species (ROS) upon pathogen encounter (Wynn et al., 2013). ROS are highly reactive signaling molecules fundamental for macrophages to obliterate invasive microorganisms (Canton et al., 2021). However, overabundant levels of ROS provoke the oxidative alteration of proteins and other biomolecules, which can cause cell death and an uncontrolled immune response (H Sies, 1985; Ueda et al., 2002). Polyunsaturated fatty acids, which are major constituents of cellular membranes, are especially vulnerable to such oxidate alterations, resulting in the formation of reactive aldehydes like malondialdehyde (MDA) and 4-hydroxynonenal (4-HNE). 4-HNE is abundantly produced in activated macrophages. Under normal physiological conditions, the concentration of 4-HNE rages in human blood and serum from 0.05 μM to 15 μM in human blood and serum (Dwivedi et al., 2007; Parola et al., 1999). In case of inflammation, the 4-HNE concentration can exceed 100 μM, which has a toxic effect on microbial and host cells and stimulates apoptosis (Smathers et al., 2012).

4-HNE is not a stable metabolite. 4-HNE’s chemical structure contains three reactive groups, including a double bond, a carbonyl group, and a hydroxyl group, which makes 4-HNE highly reactive towards nucleophilic thiol and amino groups (Dalleau et al., 2013). This allows 4-HNE to react with many proteins, resulting in the accumulation of 4-HNE protein adducts (Ayala et al., 2014; Schaur, 2003). As signaling proteins can be covalently modified with 4-HNE, this can alter cellular functions (Ayala et al., 2014).

For this reason, the treatment of immune cells with 4-HNE modifies the immune response. For instance, elevated concentrations of 4-HNE (25 μM to 50 μM) stimulate COX-2 mRNA levels and trigger the activation of the p38 MAPK signaling pathway in RAW264.7 murine macrophages (Kumagai et al., 2002, 2004). In J774 murine macrophages, exposure to 4-HNE at concentrations ranging from 0.1 μM to 10 μM elevates TGF-β mRNA levels (Leonarduzzi et al., 1997). In the human monocytic cell line THP-1, 4-HNE inhibits NF-κB activation dose-dependently by blocking the phosphorylation of the inhibitory protein IκBα (Page et al., 1999). Our previous work revealed that in human monocyte-derived macrophages, 50 μM of 4-HNE selectively blocked the production of IL-10 by inhibiting the NF-κB pathway (Melina Ioannidis, 2025). Moreover, transcriptomics analysis revealed that 4-HNE provoked pronounced transcriptional reorganization in the macrophages.

Surprisingly, we found that the gene that was most upregulated consistently in all 8 tested donors was a neuronal protein that is not detectable in the absence of 4-HNE: Synaptotagmin 1 (SYT1). SYT1 is one of the most highly studied membrane proteins because it is the primary calcium sensor for neurotransmitter secretion (Chapman, 2002). It is expressed at very high levels in neurons and neuroendocrine cells, where SYT1 facilitates rapid neurotransmitter release into the neuronal synapse upon Ca^2+^ influx. SYT1 senses Ca^2+^ through two C2 domains, thereby triggering membrane fusion of neurotransmitter-carrying vesicles with the plasma membrane. This results in neurotransmitter release into the synaptic cleft (Chapman, 2002). Given its involvement in neurotransmitter release, it is not surprising that SYT1 is exclusively expressed in neurons and neuroendocrine cells (Bradberry et al., 2020).

It is unclear why 4-HNE stimulated macrophages to express SYT1, and there is no known role of SYT1 outside the neuronal system. However, other members of the synaptotagmin family are known to affect phagocytosis in immune cells, namely SYT5, SYT7, and SYT11. SYT5 is a Ca^2+^-sensing SYT form detected in whole cell- and phagosome lysates of the murine macrophage cell lines RAW264.7 and J774. In peritoneal exudate macrophages obtained from female C57BL/6 mice, SYT5 co-localizes with the recycling endosome marker transferrin receptor and early endosomal antigen (EEA) 1. The silencing of SYT5 with siRNA in RAW2647 cells led to an impairment of phagocytosis (Vinet et al., 2008). Mouse bone marrow-derived macrophages express SYT7, another Ca^2+^ sensing member of the SYT family. The deletion of SYT7 has been shown to limit the ability to ingest zymosan particles in bone marrow-derived macrophages, showing the necessity of SYT7 for phagocytosis (Becker et al., 2009). Finally, bone marrow-derived and RAW264.7 murine macrophages express SYT11, which is localized on phagosomes and lysosomes (LAMP-1), and the siRNA knockdown of SYT11 increases cytokine secretion and phagocytosis (Arango Duque et al., 2013).

Based on these findings, we hypothesized that SYT1 would be involved in phagocytosis. By overexpressing SYT1 fused to a fluorescent reporter protein, we observed that SYT1 predominantly localizes on the plasma membrane in macrophages. Building on this finding, along with the known roles of other SYT isoforms in macrophages and related cell types, we investigated SYT1’s involvement in phagocytosis. Functional assays confirmed that both 4-HNE and SYT1 inhibit the phagocytosis of pathogenic bacteria by human macrophages. These results uncover an unexpected role for SYT1 in immune cells.

## Methods

### Isolation CD14+ monocytes and macrophage differentiation

Buffy coats of healthy donors were obtained from the Dutch blood bank. The Dutch Blood Bank approved the utilization of human blood, adhering to national and institutional guidelines during all experiments. The Dutch blood bank obtained informed consent from all donors. The investigators could not determine the identity of the blood donors as the samples were anonymized.

A standard density gradient centrifugation Lymphoprep (STEMCELL Technologies, 07861) was used to extract peripheral blood mononuclear cells (PBMC) from buffy coats as described earlier (Melina Ioannidis, 2025b). Subsequently, magnetic-activated cell sorting (MACS) using CD14 microbeads (Miltenyi Biotec, 130-114-976) was used to separate monocytes from the PBMC fraction. The CD14+ monocytes were cultured in ultra-low adherent 6-well plates (Corning, CLS3471-EA) for seven days with 2 mL of complete media containing RPMI 1640 (Gibco, 11530586), 10% fetal bovine serum (FBS) (Hyclone, 10309433), 2 mM L-glutamine (Gibco, 15430614), 1% Antibiotic-Antimycotic (Gibco, 15240062) supplemented with 100 ng/mL recombinant human M-CSF protein (R&D systems, 216-MC) at 37 °C and 5% CO2. After 7 days of differentiation, macrophages were washed with room temperature PBS and incubated with cold PBS at 4°C for 30 min to detach macrophages. Macrophages were pre-treated with different concentrations of 4-HNE (Sigma/MERCK, 393204) for 10 min at 37°C in complete RPMI. Afterward, cells were stimulated with 100 ng/mL LPS (Sigma, L4391-1MG) for different time points at 37°C.

### RNA sequencing: Experimental procedure

Naive macrophages were treated with 0 μM or 50 μM 4-HNE prior to LPS (100 ng/mL) stimulation for 24h. Thereafter, cells were washed thrice with cold PBS before lysing the cells with 1 thioglycerol/homogenization solution. The RNA was isolated and sequenced by the RNA facility at University Medical Center Groningen, the Netherlands.

### RNA sequencing: Read mapping and differential expression analysis

RNA-seq data was aligned to the hg38 assembly of the human genome using STAR v2.7.10b (Dobin et al., 2013)(parameters: --outFilterMultimapNmax 20 --alignSJoverhangMin 8 --alignSJDBoverhangMin 1 --outFilterMismatchNmax 999 –outFilterMismatchNoverReadLmax 0.04 --alignIntronMin 20 --alignIntronMax 1000000 --alignMatesGapMax 1000000). FeatureCounts v2.0.0 (Liao et al., 2014) and the latest refFlat annotation from UCSC Gene counts (parameters: -M -p) were used to compute the gene counts. Differential expression analysis was performed using DESeq2 v1.40.1. (Love et al., 2014). Pathway enrichment analysis was performed using the package pathfindR v2.2.0. (Ulgen et al., 2019).

### Reverse transcription-quantitative polymerase chain reaction (RT-qPCR)

To confirm the RNA-seq data for selected genes, naive macrophages were treated with 0 μM, 0.5 μM, or 50 μM 4-HNE for 10 min at 37°C before LPS stimulation. Samples were assembled after 6 hrs and 24 hrs LPS stimulation. Thereafter, the Quick-RNA MiniPrep kit (ZymoResearch, R1054) was used to isolate RNA following the manufacturer’s instructions. Isolated RNA was used to produce cDNA using the Moloney murine leukemia virus reverses transcriptase (M-MLV-RT) kit (Invitrogen, 28025-021) following manufacturer guidelines. cDNA (2 ng/μL per reaction) was used for qPCR using 5 μL SYBR Green Master Mix (Applied Biosystems, A25742) and 2 μL primer mix (10 μM FwD + 10 μM RV) as displayed in Table 1. qPCR was performed using the StepOnePlus Real-Time PCR System (ThermoFisher) using the following protocol: 50°C for 2 min, 95°C for 2 min, 95°C for 15 s, and 60°C for 1 min. This protocol was repeated 40 times. The results were analyzed using the 2-ΔΔCt method as described earlier (Livak & Schmittgen, 2001).

**Table 1.**
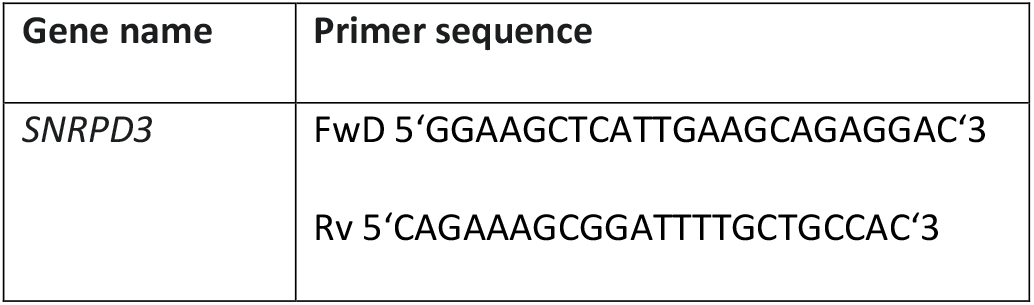
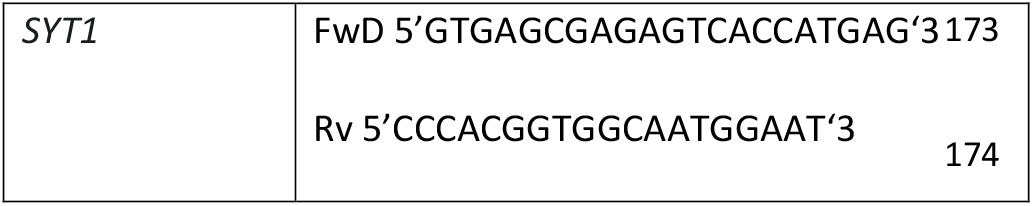
Primers for RT-qPCR. All primers for human. All primers were from Sigma Aldrich.

### Labelling of Zymosan particles

Zymosan A from Saccharomyces cerevisiae (Sigma Aldrich, 58856-93-2) were incubated 30 min on a shaker with Atto647N NHS esters (Sigma Aldrich, 18373) or Alexa Fluor 633 NHS Ester (Succinimidyl Ester) (Invitrogen, A20005). Subsequently particles were washed three times in PBS and stored at −20°C.

### Transfection of macrophages with SYT1-GFP and confocal imaging

To transfect macrophages the Neon transfection system (Fisher Scientific, 10090314) with the 10 μl tip protocol (1000 V, 40 ms. 2 pulses) was used. Using 2 μg plasmid per transfection of 1.0 x 10^6^ cells with p156rrl-pCMV-pH(ecliptic GFP)-TEV-rnSyt1 construct (Tawfik et al., 2021), kindly gifted from Jakob Balslev Sørensen, in T buffer. Cell were transferred to ultra-low adherent 6-well plates (Corning, CLS3471-EA) with warm phenol red-free RPMI (Gibco, 32404014), supplemented with L-glutamine, and incubated for 1.5h at 37°C before adding 10% FBS and 100 ng/µL M-CSF. The next day, macrophages were washed with room temperature PBS and incubated with cold PBS at 4°C for 30 min to detach before seeding them in a 4-chamber microscopy dish (Greiner, 627870) with the addition of 30 Zymosan particles per cell, labeled with Atto647N NHS esters or Alexa Fluor 633 NHS Ester as described above, per cell. The cells were incubated for 2-4 hrs at 37 °C cells. The last 30 min cells were incubated with DPPE-Abberior STAR 580 membrane dye (1 μg/mL, Abberior, ST580) and were imaged directly or after fixation with 4% PFA, using an LSM800 Zeiss microscope with 63x oil immersion lens.

Fixed cells were washed three times with PBS and subsequently blocked and permeabilized for 30 min at 4°C using CLSM buffer containing PBS, 20 mM glycine (BoomBv/Acros Organics, 220910010), 3% BSA (Merck, 126575), and 0.1% saponin (Sigma, 47036-50G). Thereafter, cells were stained overnight at 4°C with a primary antibody for SYT1 (SYSY Antibodies, 105011, 1:500), LAMP1 (Sigma, L1418,1:140), VAMP3 (Novus Biological, NB300-510, 1:100) in CLSM with 0.1% saponin (Sigma, 47036-50G) at 4°C. The next day, the cells were washed three times with PBS and 0.1% saponin (Sigma, 47036-50G) and incubated with secondary antibody donkey-anti mouse IgG (H&L) Alexa fluor 488 (ThermoScientific, 10544773, 1:400) and donkey-anti rabbit IgG (H&L) Alexa fluor 568 (ThermoScientific, 10544773, 1:400) for 30 min at room temperature in CLSM with 0.1% saponin (Sigma, 47036-50G). Subsequently, cells were washed three times with PBS and 0.1% saponin (Sigma, 47036-50G), and the coverslips were mounted on glass slides in 67% glycerol containing 1 mM Trolox (Sigma Aldrich, 238813) and 0.33 μg/ml DAPI (Sigma Aldrich, 32670). Samples were imaged using an LSM800 Zeiss microscope with a 63× 1.4 NA oil lens. Image analysis was performed in FIJI with custom macros (Supplementary Figure 1A & B). The Pearson correlation coefficient was calculated using the JACoP plugin (BOLTE & CORDELIÈRES, 2006).

### Protein expression and phosphorylation by western blot

For western blot, 1,000,000 macrophages were seeded in a 6-well plate and treated with or without 50 μM 4-HNE prior to LPS stimulation, as described above. Thereafter, the cells were washed thrice with cold PBS before lysing them with RIPA buffer (Thermo Scientific, 89900) containing Complete Mini EDTA-free Protease Inhibitor Cocktail (Sigma, 11697498001) and phosSTOP (Sigma/Merck, 4906837001) following manufacturer guidelines. The samples were denatured and unfolded using 4× Laemmli sample buffer (Bio-Rad, 1610747), heated at 90°C for 5 min, and separated using a 4-20% Mini-PROTEAN TGX Precast protein gels, 10 well, 50 μl (Bio-Rad, 456-1096) at 100 V. Proteins were blotted onto poly(vinylidene difluoride) (PVDF) membranes (Bio-Rad, 1620177) for 60 min at 90 V. Subsequently, membranes were washed three times in UltraPure Water before blocking with Intercept Blocking Buffer (LiCor, 927-60001) for 60 min at RT. After blocking, blots were incubated with primary antibody (Table 2) overnight at 4°C. The next day blots were washed three times and incubated for 60 min with secondary antibody at RT. Subsequently, membranes were incubated with anti-rabbit IgG, HRP-linked Antibody (Table 2) treated with WesternSure PREMIUM Chemiluminescent Substrate (LiCor, 926-95000) before imaging using the Li-Cor Odyssey Fc Imaging system and analyzed using Image Studio Software.

**Table 2.** Antibodies used in this study.

| Antibody | Company | Catalogue number | Dilution |
| --- | --- | --- | --- |
| Synaptotagmin1<br>antibody<br>cytoplasmic tail | SYSY Antibodies | 105011 | 1:500 |
| GAPDH (14C10)<br>Rabbit mAb | Cell Signaling<br>Technologies | 2118 | 1:1000 |
| Anti-rabbit IgG, HRP-linked Antibody | Cell Signaling<br>Technologies | 7074S | 1:3000 |
| Anti-mouse IgG,<br>HRP-linked Antibody | Thermo fisher | 31430 | 1:5000 |

### Isolation and generation of mouse bone-marrow-derived macrophages

The tibia and femur were isolated to generate bone-marrow-derived macrophages (BMDM) from female and male WT C57BL/6J and SYT Hz C57BL/6J mice. Bones were gifted by Prof. Dr. Matthijs Verhagen from the Neurosciene-Cellular & Molecular Mechanism Department at the Vrije Universiteit Amsterdam. Pre-mortem the animals were housed, bred, and handled in accordance with Dutch and European Union (EU) governmental guidelines. The protocols were approved by the Vrije Universiteit Animal Ethics and Welfare Committee (approval number FGA 11-06) (Díez-Arazola et al., 2020).

The bones were flushed out with BMDM media (RPMI 1640 (Gibco, 11530586), 10% FBS, 2 mM L-glutamine, 1% Antibiotic-Antimycotic, and 55 µM beta-mercaptoethanol (Gibco, 31350010)) to obtain bone marrow cells. Bone marrow cells were cultured at 2 – 4 x 106 cells per Petri dish in 13 mL BMDM media containing 50 ng/mL recombinant murine M-CSF (Preprotech, 315-02) for 7 days.

### Phagocytosis of *Staphylococcus aureus*

Macrophages were 10 min pre-treated with 4-HNE before LPS (100 ng/mL) stimulation for 24h at 37°C. Thereafter, cells were treated with 5 particles per cell of *Staphylococcus aureus* (Wood strain without protein A) BioParticles, Alexa Fluor 488 conjugate (ThermoFisher, S23371/10093342) for 30 min at 37°C and particle uptake was measured using the CytoFlex S (Beckman Coulter). The data were analyzed using NovoExpress Software (Agilent).

## Results

### 4-HNE induces SYT1 expression in macrophages

To determine the effect of 4-HNE on human monocyte-derived macrophages, we re-analyzed transcriptomics data from our previous study (Melina Ioannidis, 2025), where cells were treated with LPS in the presence and absence of 4-HNE for 24h. RNA sequencing revealed that *SYT1* was the highest upregulated gene when treated with 4-HNE (Figure 1A). Even with the large variation among different blood donors, the upregulation was significant between LPS versus LPS + 4-HNE (Figure 1B). The upregulation of SYT1 was confirmed by qPCR for 6 hrs and 24 hrs, whereas the 6 hrs treatment showed a higher increase compared to 24 hrs in the 4-HNE treated condition (Figure 1C). To validate the expression of SYT1 at the protein level, the effect of 4-HNE on SYT1 expression was tested by western blot for three donors. Human monocyte-derived macrophages were pre-treated with 4-HNE for 10 min, followed by LPS stimulation for 24 hrs. The western blot showed upregulation of SYT1 in cells treated with 4-HNE in the presence and absence of LPS (Figure 1D & E).

**Figure 1.**
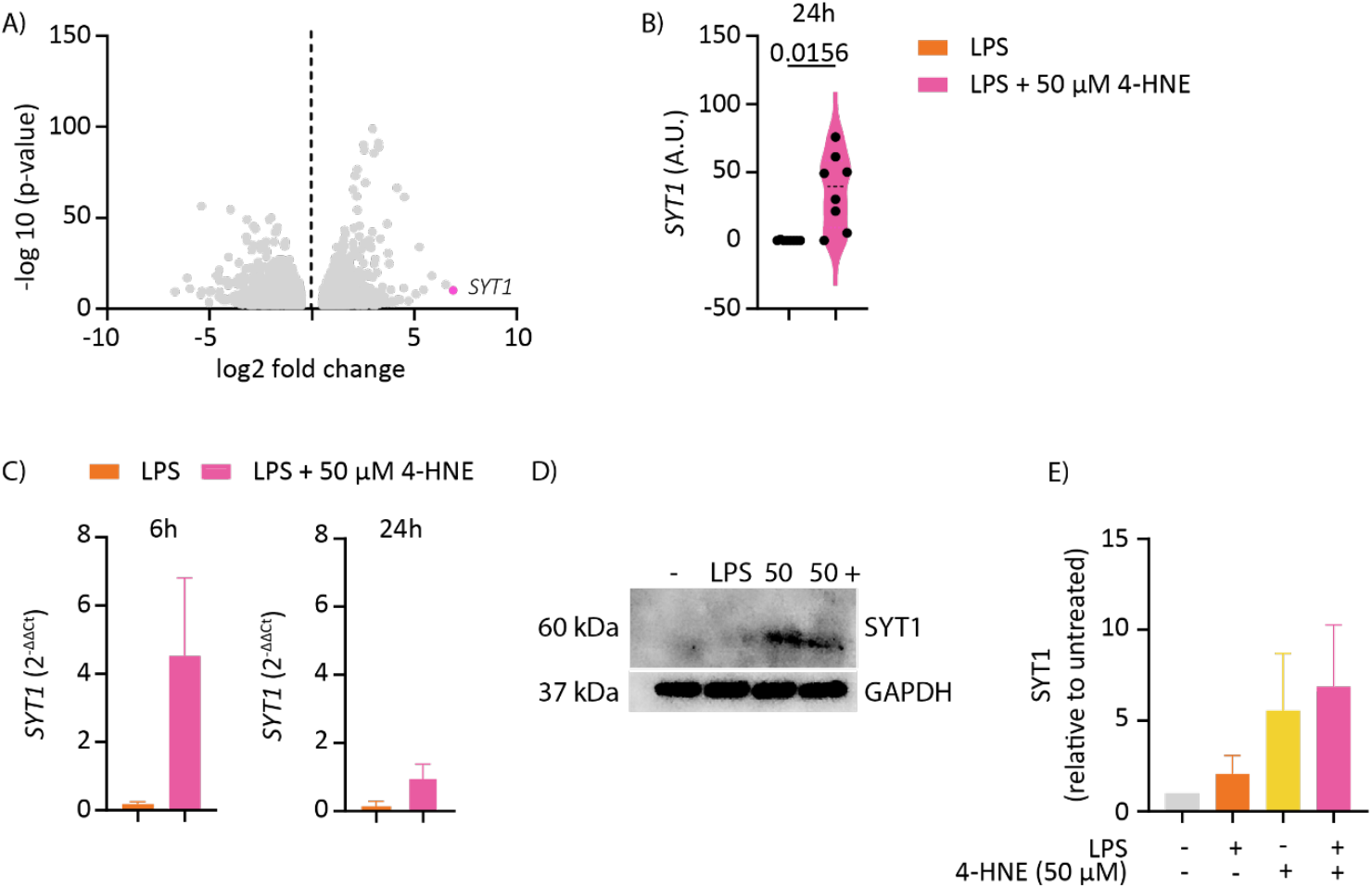
4-HNE induces the expression of SYT1 in macrophages. Human peripheral blood monocyte-derived macrophages were treated with and without 4-HNE in the presence of LPS for 24 hrs. A) Volcano plot of RNA sequencing data shows the most upregulated gene in the presence of 4-HNE is *SYT1* (pink). Each data point represents one gene; data are for n=8 donors. B) The violin plot shows the expression for *SYT1*. Each data point represents one donor; data are tested for normality with the Sapiro-Wilk normality test, analyzed using a paired t-test, and the p-value is displayed. C) RT-qPCR confirmed RNA sequencing data for 6 hrs and 24 hrs of treatment (n=3 donors). Data are displayed as 2^-ΔΔCt^ and plotted as mean with SEM. D) Representative Western blot for SYT1 (60 kDa) and GAPDH (37 kDa). E) Quantification of western blot results for n=3 donors, displayed relative to untreated and as mean value with SEM.

### SYT1-GFP localizes at the plasma membrane

We next investigated where SYT1 is localized in macrophages. We used overexpression of SYT1 fused to GFP (SYT1-GFP) because the expression levels of SYT1 in 4-HNE treated macrophages varied strongly among different blood donors, and expression levels overall were too low for immunofluorescence labeling. The macrophages were transfected with p156rrl-pCMV-pH(ecliptic GFP)-TEV-rnSyt (SYT1-GFP) (Tawfik et al., 2021). SYT1-GFP was located at the plasma membrane in both live and fixed cells (Figure 2A).

**Figure 2.**
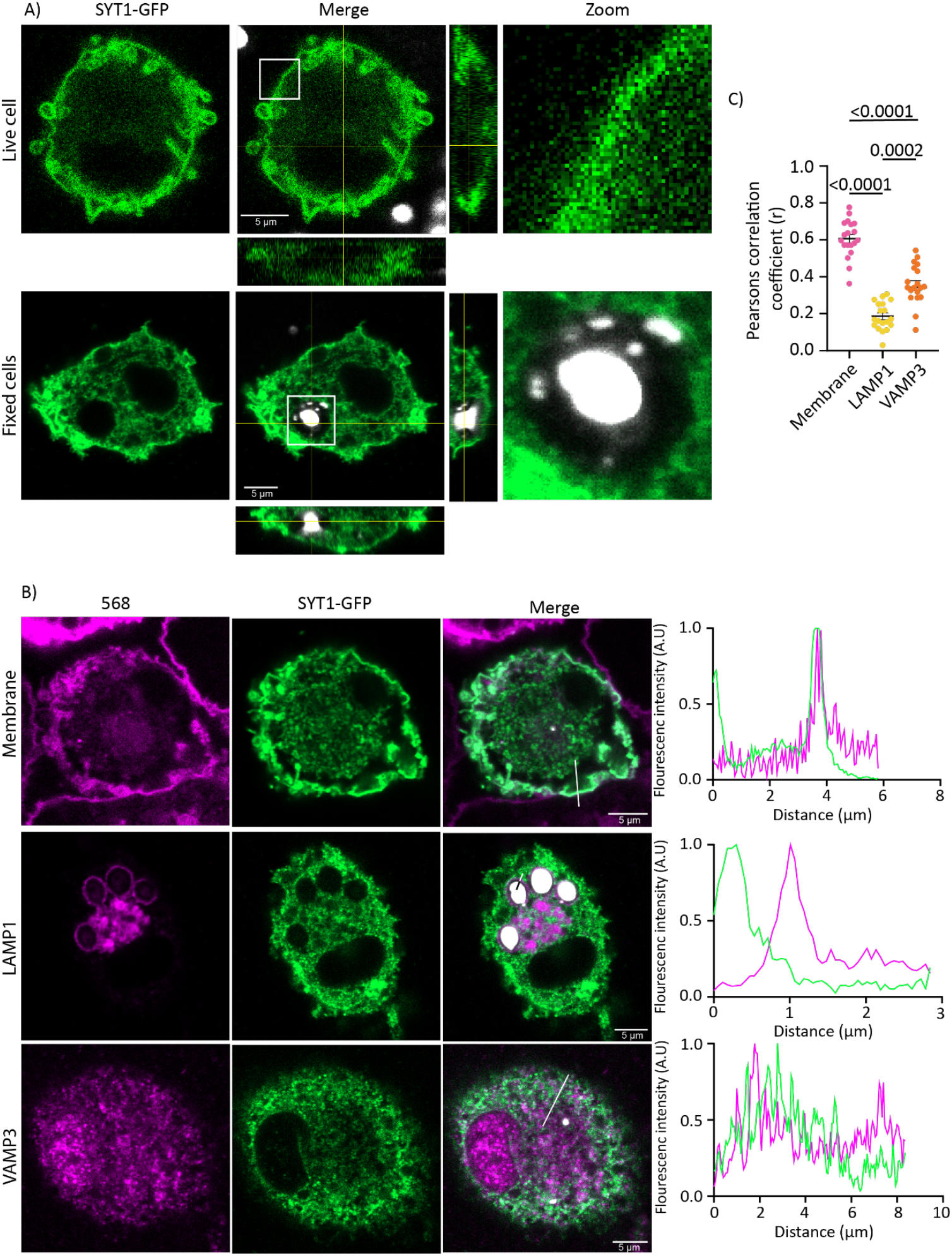
SYT1-GFP localizes at the plasma membrane. Human peripheral blood monocyte-derived macrophages were transfected with SYT1-GFP (green) and treated with zymosan labeled with Alexa fluor 647 (white). A) Representative confocal images of a live cell (upper row) and a fixed cell with a phagosome (lower row). The orthogonal views are shown for the merge. The scale bar represents 5 microns. B) Representative confocal images of fixed cells expressing SYT1-GFP (green) and incubated with zymosan Alexa fluor 647 (white; only shown in the merge). The cells were labeled with an Alexa fluor 568 conjugated membrane probe, immunolabeling for lysosomal marker LAMP1 or endosomal marker VAMP3 (magenta). The scale bars indicate 5 microns, and the lines indicate the positions of the fluorescence intensity profiles (right panels). C) The Pearson correlation coefficients were determined for the membrane probe, LAMP1, and VAMP3 for 3 donors. Data are represented as median, and each data point represents one cell. Data were tested for normality using the Sapiro-Wilk test, and one-way ANOVA was used to test for significance. The p-values are displayed.

As previous research has shown that SYT5, SYT7, and SYT11 influence phagocytic capacity in macrophages (Arango Duque et al., 2013; Becker et al., 2009; Vinet et al., 2008), the macrophages overexpressing SYT1-GFP were also treated with Zymosan Atto647 particles, a common phagocytosis model cargo. However, we did not detect SYT1-GFP at the phagosomal membrane. To further dissect the location of SYT1-GFP, we co-stained for different organelles using the DPPE-Abberior STAR 580 membrane dye, the lysosomal-associated membrane protein (LAMP) 1, and vesicle-associated membrane protein (VAMP) 3. Z-stack images were acquired from samples of three different donors, and the co-localization was quantified using the Pearson correlation coefficient. LAMP1 did not correlate with SYT1-GFP, whereas there was some overlap of SYT1-GFP with VAMP3. However, the membrane dye showed the most overlap with SYT-GFP (Figure 2B-C). Thus, at least overexpressed SYT1-GFP localizes at the plasma membrane in macrophages.

### SYT1 inhibits phagocytosis in macrophages

As we observed markedly less particle uptake in our SYT1-GFP transfected cells by microscopy, and previous studies showed that other SYT forms influence phagocytic capacity in macrophages (Arango Duque et al., 2013; Becker et al., 2009; Vinet et al., 2008), we investigated the effect of SYT1 on phagocytosis. First, human macrophages were treated with or without 4-HNE in the presence of LPS for 24 hrs before adding *Staphylococcus aureus* conjugated to Alexa Fluor 488 (AF488). Flow cytometry analysis showed significant downregulation of phagocytosis when treated with 4-HNE (Figure 3A), indicating that 4-HNE treatment inhibits phagocytosis.

**Figure 3.**
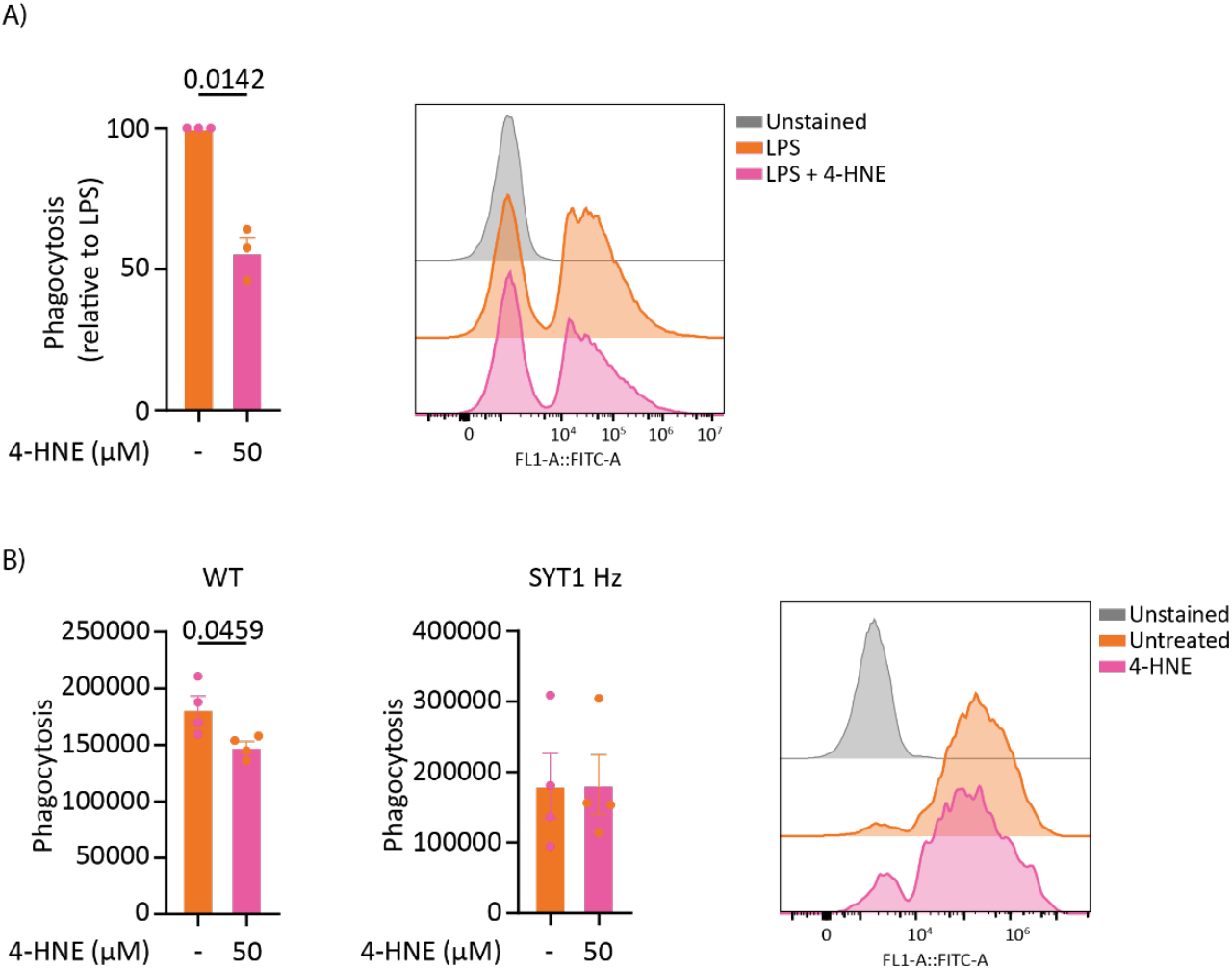
4-HNE inhibits phagocytosis in macrophages potentially through SYT1. A) Phagocytosis of human monocyte-derived macrophages (n=3 donors). Human macrophages were treated with different concentrations of 4-HNE in the presence of LPS for 24h. Phagocytosis was tested with *Staphylococcus aureus* labeled with Alexa fluor 488. Data display the median fluorescence intensity of phagocytosed bacteria normalized to LPS. Each dot represents one donor, and the error bar displays SEM. The histogram median fluorescence intensity for unstained (grey) versus LPS (orange) and LPS + 50 μM 4-HNE (pink) for one donor. B) Bone-marrow-derived macrophages of wild-type C57BL/6 and SYT1 Hz C57BL/6 mice underwent the same treatment as the human macrophages from panel A. Data display the median fluorescence intensity of phagocytosed bacteria. Each symbol represents the mean value of n=4 different mice, and the error bar displays SEM. The histograms show unstained (grey), untreated (orange), and 50 μM 4-HNE (pink) for one WT mouse. Data in A) and B) were tested for normality using the Sapiro-Wilk test, and a paired t-test was used to test for significance. The p-value is displayed in case of significant difference.

To directly link this effect to SYT1, we derived macrophages from the bone marrow of SYT1 knockout mice. SYT1 is an essential gene, and SYT1 homozygous knockout mice die 48 hrs after birth. Therefore, we used SYT1 heterozygous knockout mice (SYT1 Hz), which have a mutation in the SYT1 gene induced through homologous recombination (Geppert et al., 1994).

The heterozygous knockout mice express lower levels of SYT1 but are phenotypically normal compared to wild-type (WT) mice. Bone-marrow-derived macrophages from WT and SYT1 Hz were pre-treated with 4-HNE in the presence and absence of LPS for 24h. Phagocytosis was again investigated by measuring *S. aureus* AF488 uptake by flow cytometry. In line with our findings with 4-HNE, WT mice show a significant decrease in phagocytosis when treated with 4-HNE. In contrast, treatment of the SYT1 Hz with 4-HNE did not show a clear effect (Figure 3B).

## Discussion

SYT1 is essential for the calcium-dependent exocytosis of neurotransmitters into the synaptic cleft and modulates synaptic vesicle endocytosis for neurotransmitter recycling. Deletion of SYT1 is embryonically lethal in mice (Geppert et al., 1994) and mutations in SYT1 can result in the neurodevelopmental disorder Baker-Gordon Syndrome in humans, showing the importance of SYT1 (Baker et al., 2018; Melland et al., 2022). SYT1 is usually expressed only in neuronal cells (Bradberry et al., 2020), so we were surprised to find that the oxidative lipid metabolite 4-HNE consistently induces SYT1 expression in human peripheral blood monocyte-derived macrophages. Under normal conditions, SYT1 is not detectable in macrophages. Upon stimulation with 4-HNE in the presence and absence of LPS, we observed an upregulation of SYT1 at both mRNA and protein levels for all donors tested.

Previous research has shown the expression of other SYT family members in macrophages, their involvement in the phagocytosis of pathogens, and exocytosis of cytokines like TNF-α and IL-6. For instance, the overexpression of SYT11 in RAW264.7 murine macrophages reduces zymosan uptake (Arango Duque et al., 2013). Another member of the SYT family, SYT5, is expressed in RAW264.7 and J774 murine macrophages. Phagosome isolation showed that SYT5 is present in total cell and phagosome lysates of J774 cells. Using siRNA to silence SYT5 in RAW264.7 macrophages reduces serum-opsonized zymosan particles’ phagocytosis (Vinet et al., 2008). In human neutrophils, SYT2 translocates to the phagosome after complement-mediated phagocytosis (Lindmark et al., 2002). Although the expression level of SYT1 in 4-HNE treated macrophages was too low for immunofluorescence microscopy, our data show that overexpressed SYT1 is localized on the plasma membrane, which is similar to the plasma membrane localization of SYT2 in neutrophils (Lindmark et al., 2002). For these reasons, we tested the effects of 4-HNE on phagocytosis.

Indeed, the phagocytosis of *S. aureus* bacteria was decreased in 4-HNE-treated macrophages. To link this finding to SYT1, we obtained bone marrow from WT and SYT1 Hz C57BL/6 mice and differentiated bone marrow stem cells into macrophages. Macrophages from WT mice showed a decrease in phagocytosis when treated with 4-HNE whereas macrophages from SYT1 Hz consistently showed a lower phagocytic capacity than WT in both LPS and non-LPS conditions. These results suggest that SYT1 inhibits phagocytosis, similar to the role of SYT11 in murine bone marrow-derived macrophages (Arango Duque et al., 2013). However, whereas we found that SYT1 locates only at the plasma membrane and not at phagosomal membranes, SYT11 also localizes with LAMP1 at phagosomes (Arango Duque et al., 2013). Our findings are reminiscent of findings in neutrophils isolated from human blood, where treatment of 4-HNE also provoked a decrease in phagocytosis of FITC-labelled *S. aureus* (Chacko et al., 2016), which might also involve SYT1.

We only detected SYT1 expression in 4-HNE-treated cells. In contrast, SYT5 and SYT11 have been described as constitutively expressed by murine macrophages (Arango Duque et al., 2013; Vinet et al., 2008), whereas we could not detect the expression of these or any other SYT forms in human macrophages. Thus, 4-HNE specifically induces SYT1 expression in macrophages, potentially regulating pathogen sampling. The fact that SYT1 is only expressed in macrophages under conditions of oxidative stress indicates a tight regulation of SYT1. It would be interesting to see how 4-HNE exactly regulates SYT1 expression and if it is cell-specific.

We recently reported that 4-HNE causes a profound inhibition of the production of IL-10 by macrophages, whereas the production of other cytokines is not significantly affected (S. M. L. B. M. den O. M. S. P. G. D. I. F. B. H. B. and G. van den B. Melina Ioannidis, 2025). SYT1 might play a role here, given that other SYT forms regulate cytokine production in macrophages and neutrophils. For instance, SYT2 is present in human neutrophils and is associated with gelatinase granules (Lindmark et al., 2002). The overexpression of SYT11 in RAW264.7 murine macrophages reduces the secretion of cytokines, including TNF-α and IL-6 secretion (Arango Duque et al., 2013). Unfortunately, we could not test the effects of SYT1 on cytokine production due to low expression levels.

For the same reason, we could not test the effect of SYT1 on chemotaxis. An RNA-mediated interference screen of a T lymphoblast cell line (SupT1) revealed the involvement of SYT7, SYT5, and SYT2 in chemotaxis (Colvin et al., 2010). Infection with a virus targeting SYT5 mRNA inhibited CXCL12 and CCL2 stimulated migration of SupT1 and monocytic leukemia cell line THP-1, respectively (Colvin et al., 2010). On the contrary, infection with a virus targeting SYT2 increased the migration of SupT1 and THP-1 (Colvin et al., 2010). In the TAM2D2 mouse T cell line, SYT3 regulates the recycling and cell surface expression of CXCR4 (Masztalerz et al., 2007).

Oxidative stress leads to an increase in ROS and an accumulation of 4-HNE. The mammalian brain is vulnerable to oxidative stress (Erecińska & Silver, 2001) due to the abundant presence of PUFAs, high energy requirements, and weak antioxidant defenses (Cobley et al., 2018). Abnormal levels of ROS are related to the pathogenesis of aging-related neurodegenerative diseases like Alzheimer’s and Parkinson’s disease (Bascuas et al., 2020; Garrido-Pascual et al., 2020; Giordano et al., 2014). It is therefore no surprise that studies have shown elevated levels of 4-HNE protein adducts in body fluid of patients with Alzheimer’s disease (De Virgilio et al., 2016; Di Domenico et al., 2017), Parkinson’s disease (De Virgilio et al., 2016; Di Domenico et al., 2017; Sardar Sinha et al., 2018), Huntington’s disease (Di Domenico et al., 2017; Lee et al., 2011), and amyotrophic lateral sclerosis (Di Domenico et al., 2017; Shibata et al., 2011). Our findings could be relevant in the pathogenesis of these diseases, especially if 4-HNE also induces SYT1 expression in microglia, the brain’s macrophages.

Moreover, although an untested hypothesis, 4-HNE might also promote SYT1 expression in neurons and neuroendocrine cells. This is potentially of interest because SYT1 expression levels have been linked to neuronal diseases. For example, SYT1 is downregulated in neurons of the nucleus basalis, cerebellum, thalamus, and hippocampus of patients with Alzheimer’s disease (Sze et al., 2000; Yoo et al., 2001). Another study also reported that SYT1 is downregulated in the hippocampus of early Alzheimer’s disease patients, although in this case, the expression was unaltered in the cortex (Sze et al., 2000).

## Acknowledgments

We thank Jakob Balslev Sørensen from the Department of Neuroscience at the University of Copenhagen in Denmark for the kind gift of the SYT1-GFP plasmid.

---

**Supplementary Figure 1.**
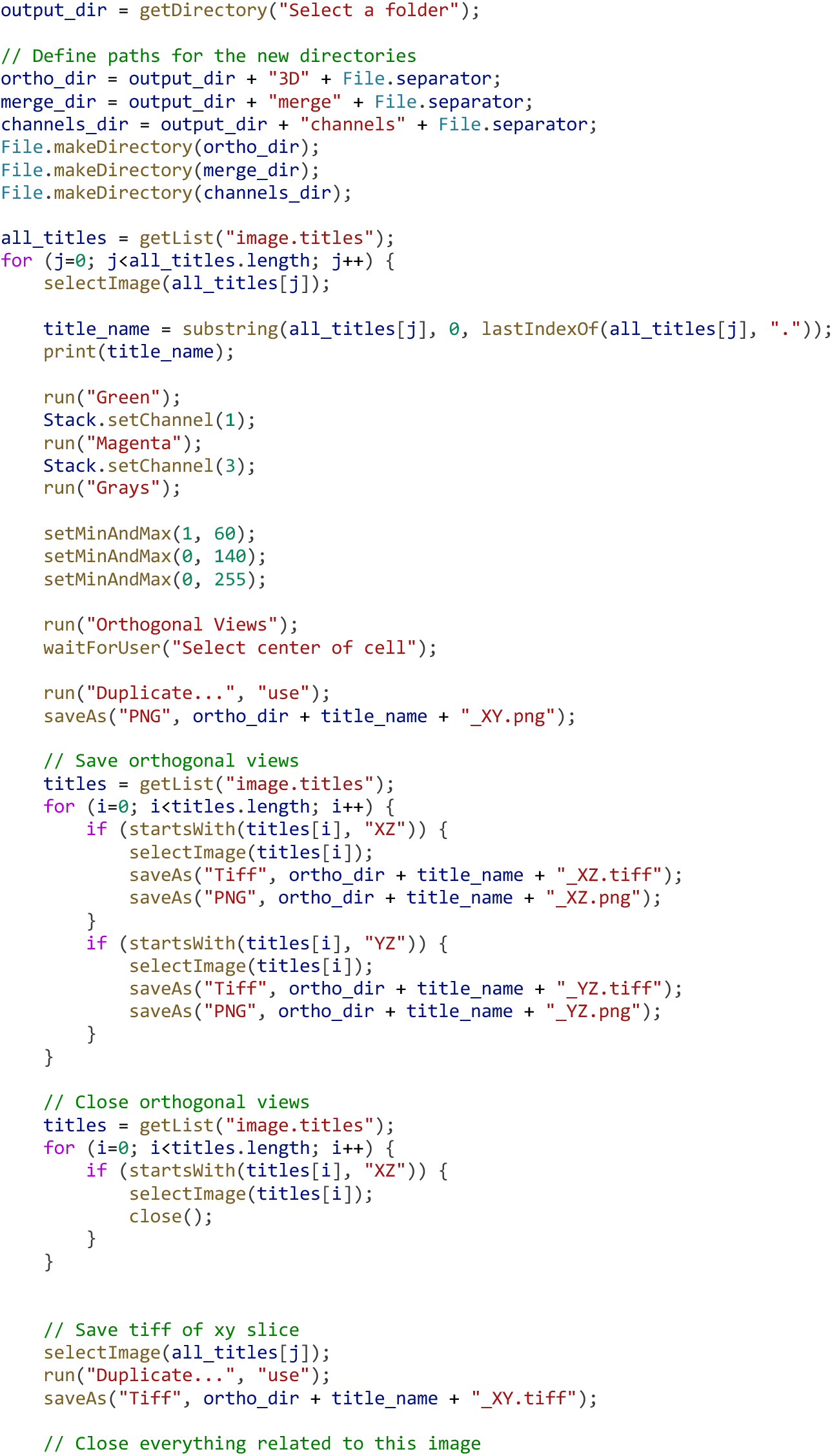
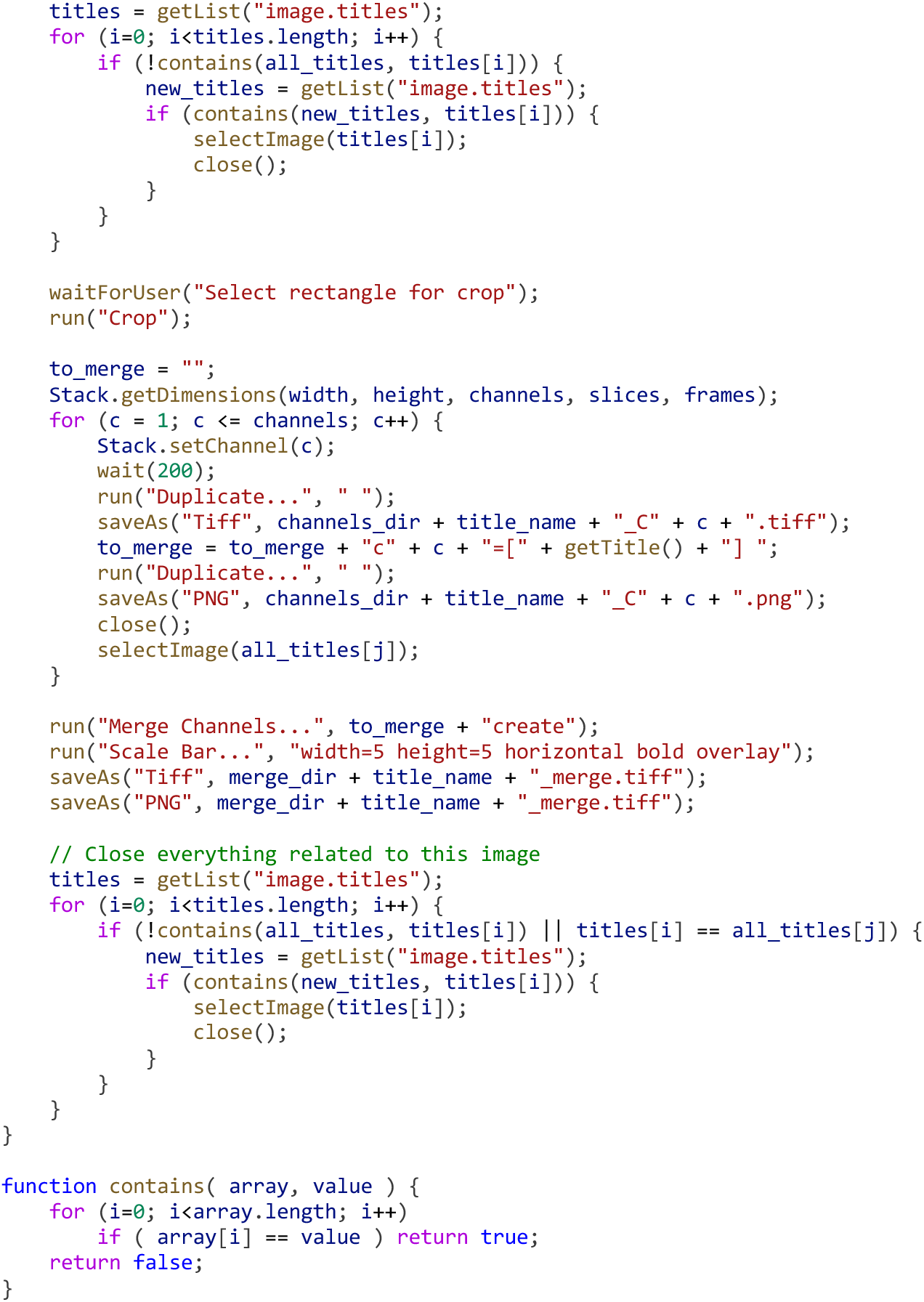
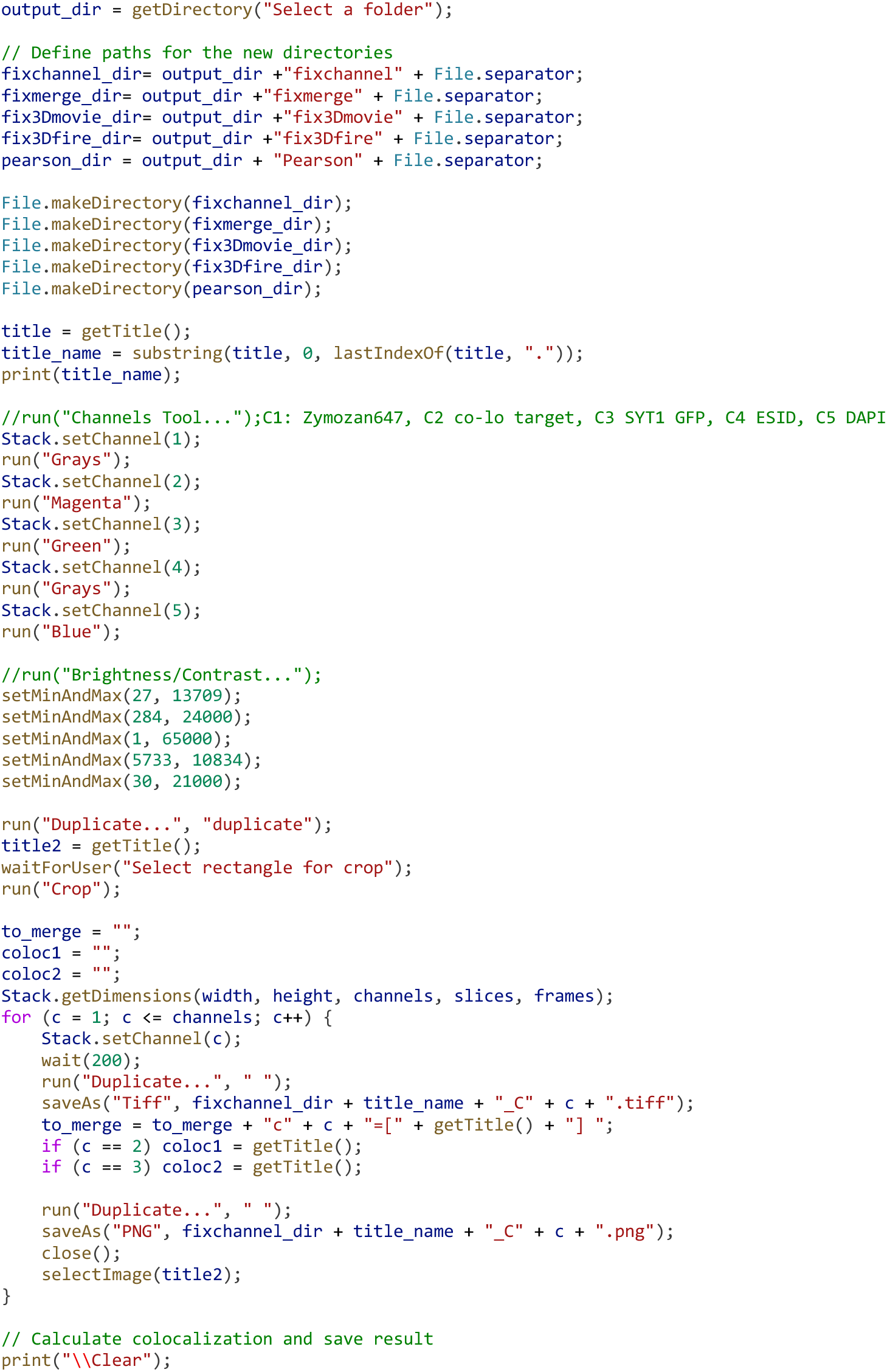
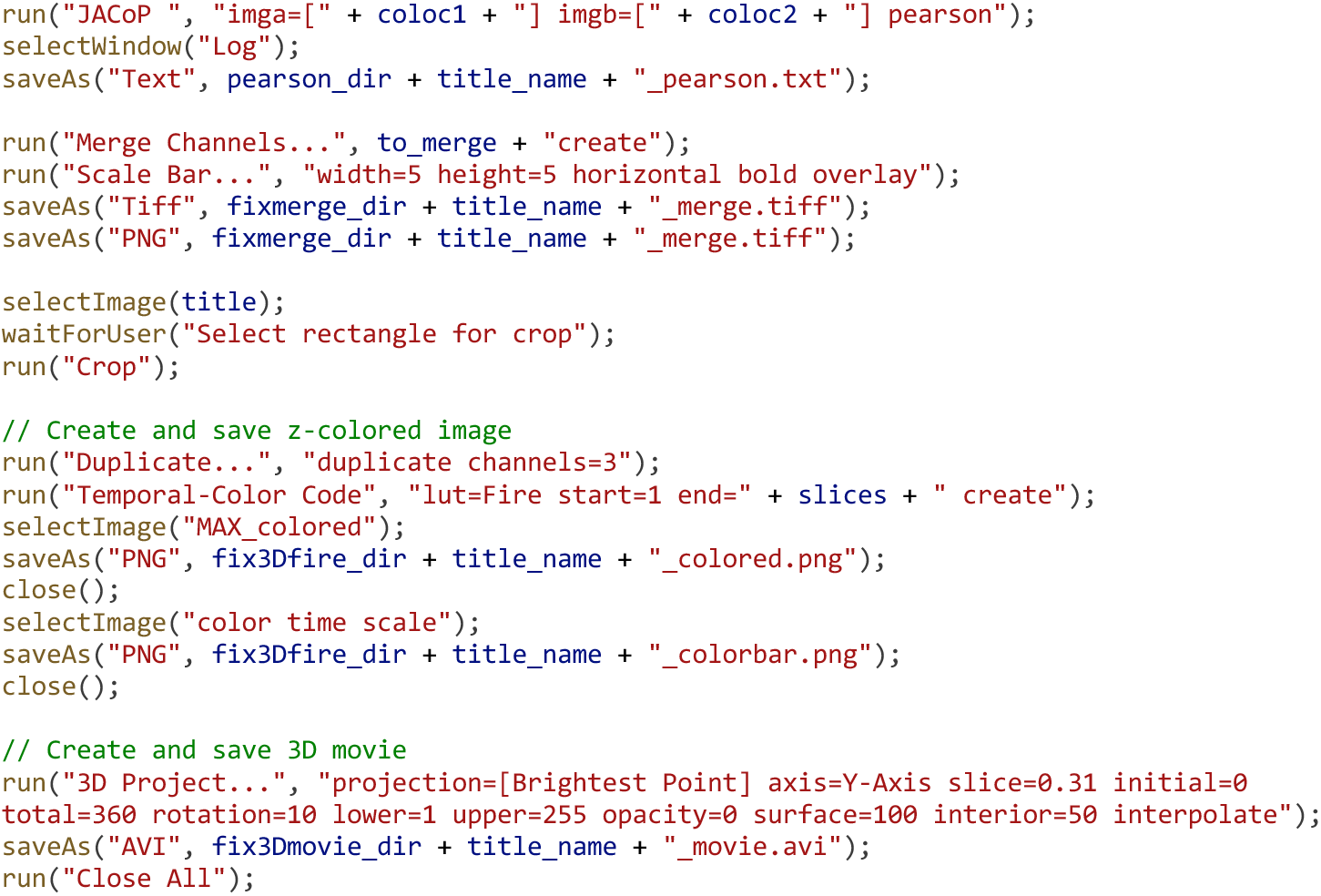
FIJI macro script used to analyze z-stack images. A) FIJI macro was used to create an orthogonal view, a single channel image, and a merge image. B) FIJI macro that was used for Pearson correlation coefficient analysis.

